# Reframing Sleep Architecture: A Compositional and Temporal Approach to Sleep Data Analysis

**DOI:** 10.1101/2025.07.04.663192

**Authors:** Mattia Bonzanni, Alice Braga

## Abstract

Significant progress has been made in developing sleep staging methodologies; however, less attention has been devoted to the analysis of sleep architecture. Two critical aspects remain underexplored: the choice of binning window (i.e., grouping data into time intervals) and the statistical treatment of the interdependencies among sleep phases. While one-hour bins are commonly used, this choice is often based on convention rather than empirical justification. Additionally, sleep architecture data are typically expressed as proportions of time spent in each sleep phase, forming compositional datasets, non-negative values that represent parts of a whole and are constrained by a constant sum (e.g., 100%). Such data violate the assumptions of traditional statistical methods, yet their compositional nature is often overlooked, compromising analytical validity. In this study, we address two key methodological challenges in sleep architecture analysis: (1) determining the optimal binning window through a data-driven approach that balances information retention and noise reduction, and (2) applying isometric log-ratio (ILR) transformation to account for the compositional structure of the data, enabling the use of conventional statistical tests. By addressing these issues, we propose a more rigorous and interpretable framework for analyzing sleep architecture, aiming to enhance the accuracy and reproducibility of findings in sleep research.

**Statement of Significance:** Sleep architecture is typically analysed without accounting for the interdependent nature of sleep stages, leading to potentially misleading interpretations. This study introduces a compositional analytical framework that treats sleep data as interdependent, where changes in one phase necessarily impact the others. By integrating compositional data principles with a biologically grounded approach to temporal binning, we offer researchers a practical and interpretable method for analysing sleep structure. This work challenges conventional analysis practices and encourages a rethinking of how sleep-related biological questions are framed. These findings hold potential relevance for improving the design and interpretation of both experimental and clinical sleep studies, ultimately supporting more accurate insights into physiological and pathological studies involving altered sleep patterns.

## Introduction

Sleep is increasingly recognized as vital for physiological and pathophysiological processes, with numerous studies investigating its multifaceted roles^1^. Sleep/wake research typically involves staging 24-hour recordings to profile behavioral states, including wakefulness, non-rapid eye movement (NREM) sleep, and rapid eye movement (REM) sleep. While substantial efforts have been dedicated to refining sleep staging methodologies, comparatively less attention has been paid to the effective presentation and analysis of sleep architecture data^2^. Sleep and wakefulness are mutually exclusive and exhaustive components, and thus any change in the duration of one state necessitates an equal and opposite change in the other. Despite this interdependence, many studies have treated sleep/wake data as independent variables, with only a few recent studies accounting for their compositional nature ^3,4^. Recognizing and appropriately analyzing the interdependent nature of sleep architecture is crucial for advancing the field and enhancing clinical applications^5,6^.

Sleep architecture data are compositional, representing parts of a whole where each component is a proportion and the sum of all parts equals a constant (e.g., 1 or 100%). This type of data is prevalent across various fields, including genomics, geochemistry, sociology, and marketing^7-11^. Compositional datasets describe the relationships between components and how they change in respect to each other^9^. However, they are incompatible with traditional statistical approaches that assume independence among variables^9,12,13^. In compositional data, components are interdependent and therefore changes in one component can only be interpreted in the context of changes in others. Moreover, compositional data exist in a constrained geometric space (simplex) rather than the usual Euclidean space, requiring specialized analytical methods. Standard visualizations like scatter plots and bar charts may not adequately capture the relationships between components due to the constant-sum constraint. Additionally, compositional datasets are sensitive to zero values, as common transformations (e.g., logarithmic) require all components to be non-zero^9,14^.

Another critical consideration in analyzing sleep/wake data is the choice of binning time. Binning serves multiple purposes, such as simplifying temporal variability to reveal trends and patterns, enhancing signal-to-noise ratio, reducing computational complexity, improving visualization, and aligning with biologically relevant time windows^15^. Common binning windows include one hour (based on zeitgeber time divisions), six hours, or light/dark cycles (12 hours each), balancing biological relevance with the need to simplify temporal trajectories. The selection of binning window size impacts the final results by altering data resolution and variability, which can affect interpretations and complicate comparisons between results derived from different bin sizes.

In this manuscript, we explore and propose strategies to address these overlooked aspects of sleep architecture analysis: determining optimal binning windows and accounting for the compositional nature of the data. We introduce appropriate visualization methods, transformation techniques, and statistical frameworks for analyzing murine sleep architecture (sleep phases: wakefulness, NREM, and REM), treating it as a valid compositional dataset.

## Material and Methods

### Binning procedure

Time-stamped summary of sleep data with phase identifiers (NREM, REM, or Wake) was used as input. If the dataset was not a continuous trace, the continuous trace was reconstructed using the initial and final times for each event. A uniform vector of binning windows was created based on the specified binning window size and, for each binning window, the proportions of each phase (NREM, REM, and Wake) were calculated. Pseudocode provided below.

#### Algorithm

Binning procedure

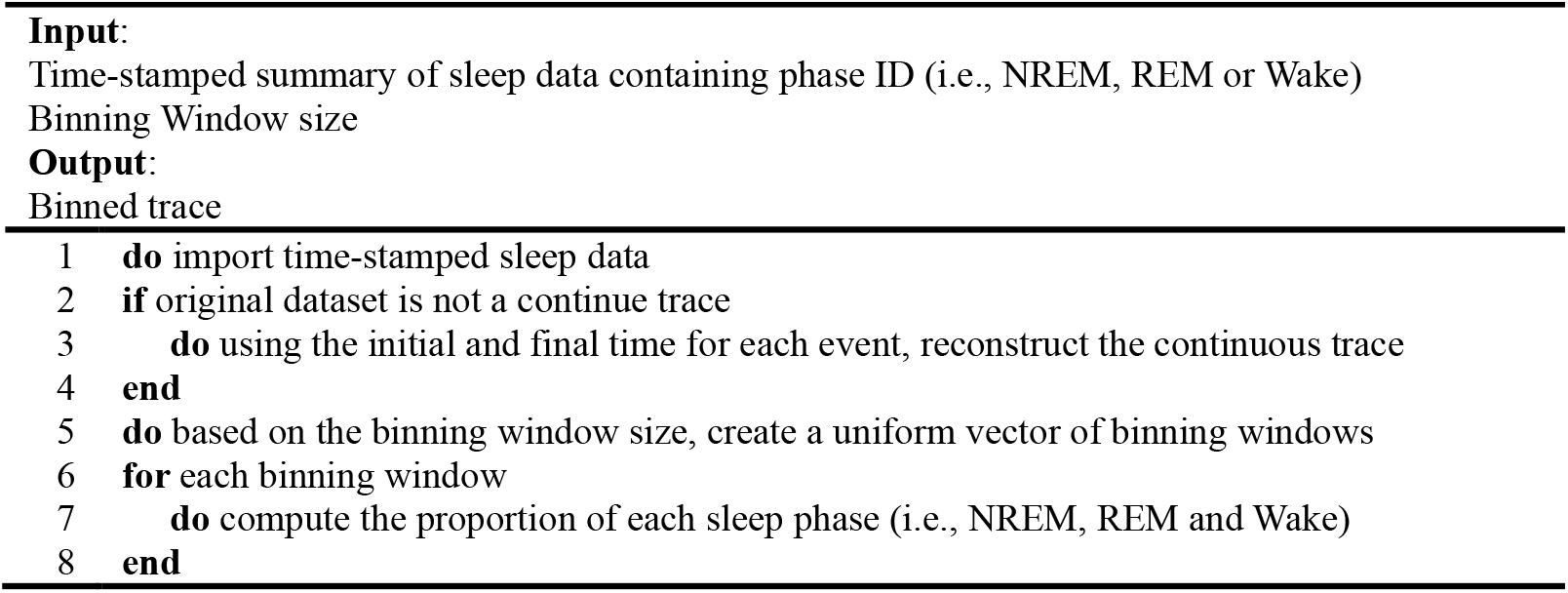

### Zero-replacement procedure

Zero elements in the binned dataset needed to be replaced (see pseudocode below). Briefly, after binning the dataset, zero elements were replaced with two-thirds of the detection limit ^9,14^. For each binning window size, the detection limit was defined as the ratio between the time resolution of the sleep staging procedure and the binning window size, effectively representing one event within the bin. In our dataset, the time resolution of the sleep staging procedure was set to 4 seconds in Sirenia. After zero-replacement, proportion were rescaled to sum to 1 (closure) to ensure compositional integrity of the dataset.

#### Algorithm

Zero-replacement procedure

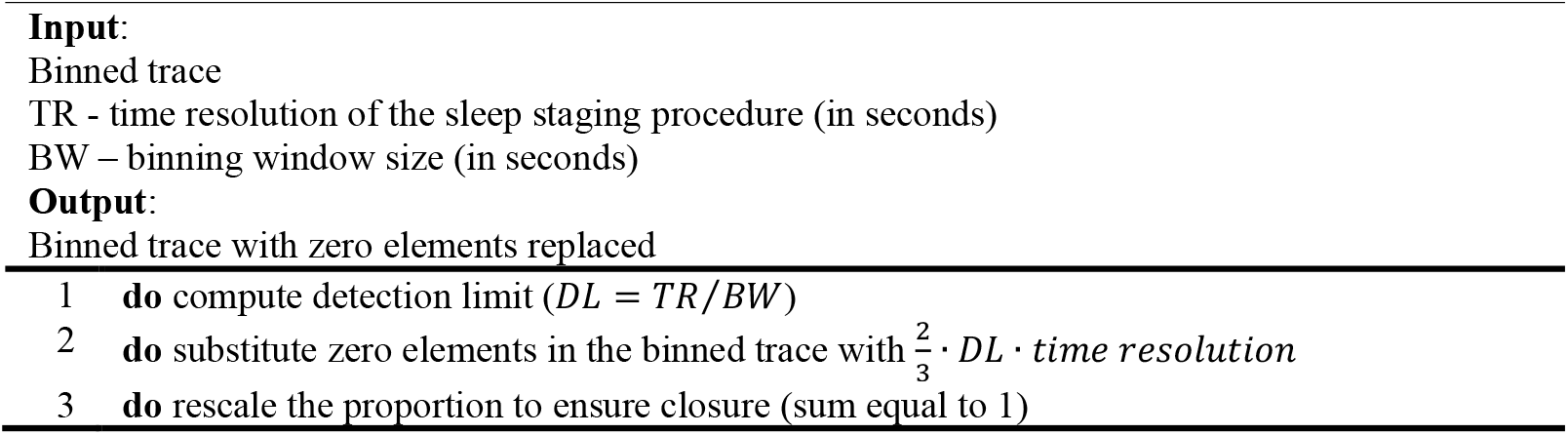

### Shannon Entropy

Shannon’s entropy of binned dataset has been computed according to:

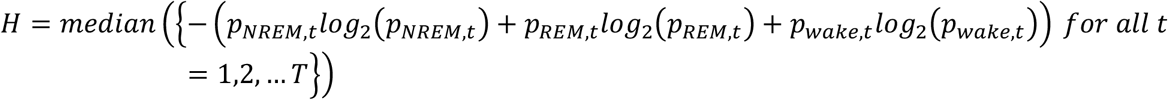

where H was the median entropy of the binned dataset, with zero elements replaced and *T* representing the total number of bins. *P*_*NREM*,*t*_, *p*_*REM*,*t*_ and *p*_*wake*,*t*_ were the probabilities of NREM, REM and wake at the *t*^*th*^ bin. Entropy was calculated for each time bin, and the median value of the resulting entropy vector was used to summarize the entropy of the entire dataset. *H* was used as a descriptive metric of the spread/uniformity of the proportions in the dataset.

### Hellinger distance

The Hellinger distance was used to compute the distortion in the original dataset caused by the zero-replacement procedure. For the dataset before zero replacement (*P*) and the dataset after zero replacement (*P*), the Hellinger distance (*d*_*H*_) has been computed for each row as follows:

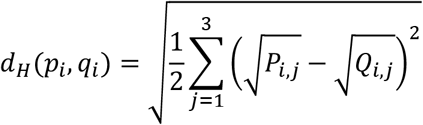

where *d*_*H*_ (*p*_*i*_,*q*_*i*_) is the Hellinger distance at the *i*^*th*^ row, and *j*^*th*^ column, *Q*_*i*,*j*_ is the value in the dataset *P* at the *i*^*th*^ row and *j*^*th*^ column. After computing the Hellinger distance for each row, rows with 0 (indicating no replacement was performed) were excluded. The median Hellinger distance of the remaining rows was then taken as a measure of distortion, focusing specifically on rows affected by the zero-replacement procedure. This approach increased the median value but provided a focused reflection of distortion regardless of the number of zero elements replaced. For a global measure of distortion, we suggest using the average Hellinger distance across all rows, as it accounts for both affected and unaffected rows, offering a more comprehensive view of the overall impact. In fact, if more than half of the rows did not have a zero element, the median would be 0. Hellinger distance values ranged in the interval [0,1], where 0 meant that two rows were identical, and 1 that two rows were maximally dissimilar. The Hellinger distance was insensitive to zero elements and scale-invariant, making it suitable for datasets with zeros. However, it did not fully capture relative changes or balance and only partially respected the compositional nature of the data. The presence of zeros in the original dataset, combined with our experimental requirements, limited the choice of techniques and prevented the direct use of the Aitchison distance, a metric specifically designed for compositional data but requiring strictly positive values.

### Isometric log-ratio transformation

The isometric log-ratio (ilr) transformation was applied to compositional data prior to statistical analysis ^9^. This transformation addressed issues of scale invariance and interdependency arising from compositional datasets. This transformation maps the data from the simplex space (constrained to a constant sum) to the Euclidean space (unconstrained). The ilr transformation expressed the composition dataset in terms of orthogonal coordinates, preserving metric properties such as distances and angles. For a given Wake-REM-NREM (W, R, N, respectively) composition, the first and second ilr coordinates (ilr_1_ and ilr_2_, respectively) were computed as follows:

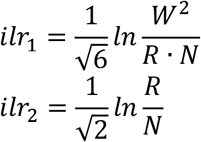

where W, R and N represented the percentages (or fraction) of each phase at a given time point (i.e., W=22%, N=65%, R=13%), *ilr*_1_ represented the proportion of wake compared to sleep (N+R) and *ilr*_2_ the proportion of REM compared to NREM.

#### Algorithm

Isometric log-ratio (ILR) transformation

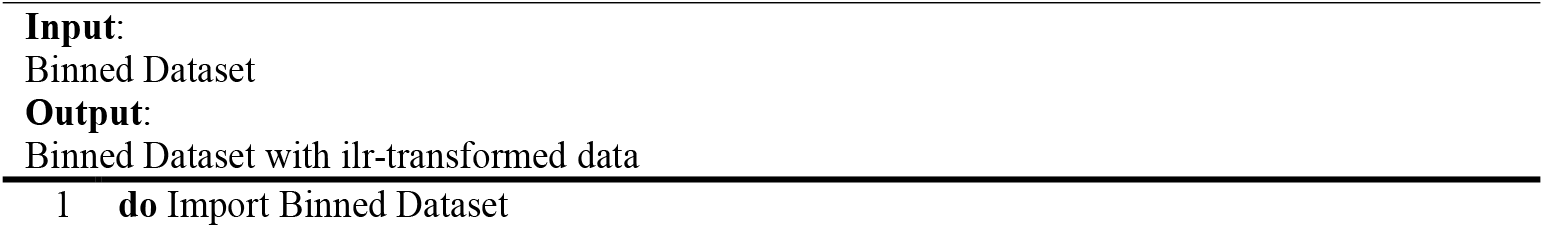
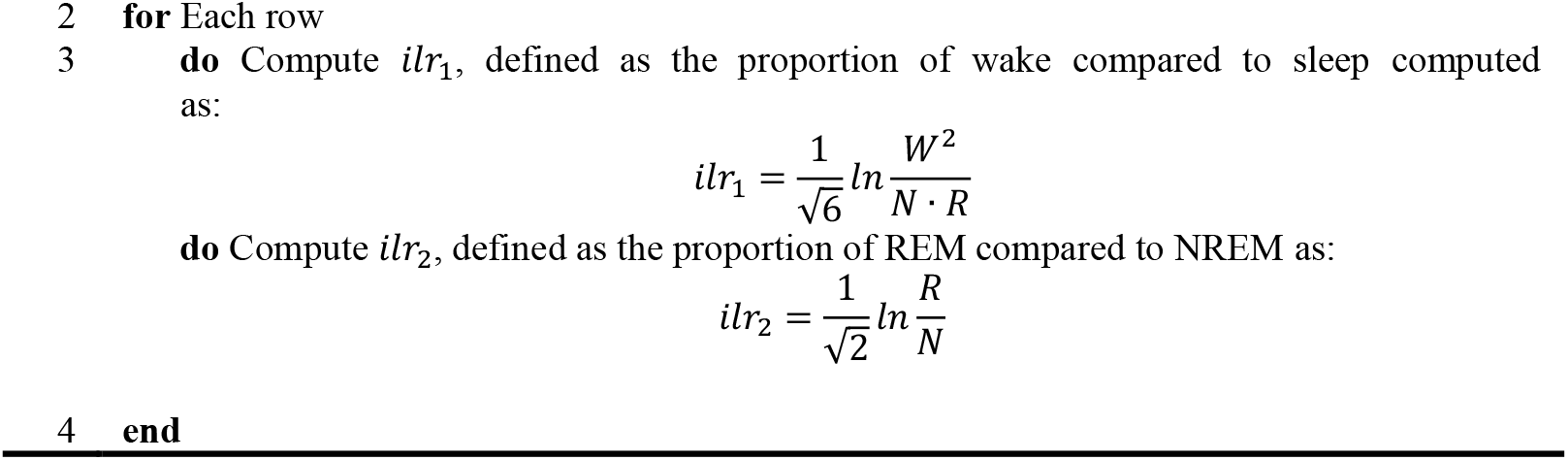

### Statistics

To address the multivariate and longitudinal nature of the dataset following isometric log-ratio transformation, repeated measures MANOVA was performed using SPSS. The computation of variates and canonical loadings was conducted using MATLAB and R by extracting information from the SSCP matrix (Sums of Squares and Cross Products), which was obtained as part of the output from the repeated measures MANOVA analysis in SPSS. The SSCP matrix served as a proxy for the variance-covariance structure, capturing the underlying variability and interdependence among the variables in the dataset. A detailed step-by-step guide for these procedures were provided in the supplementary material.

## Results

### Selection of the optimal binning window

Although binning is a common preprocessing step in sleep data analysis, there is currently no consensus or standardized guideline for selecting an appropriate binning window size. In this study, we implemented a data-driven approach as an alternative to biologically predefined binning intervals. To test this method, we used a previously published dataset (n = 23)^16^ and here we report some examples. Using an information theory framework, we sought to optimize the entropic content of the binned dataset, balancing information retention with noise reduction. Additionally, we evaluated the impact of zero elements (bins in which one or more sleep stages were not detected) on the accuracy of compositional analysis, as zero-replacement is a necessary step in handling compositional data (Fig. 1A). The optimal binning window should therefore minimize also the distortion introduced during zero-replacement.

**Figure 1:**
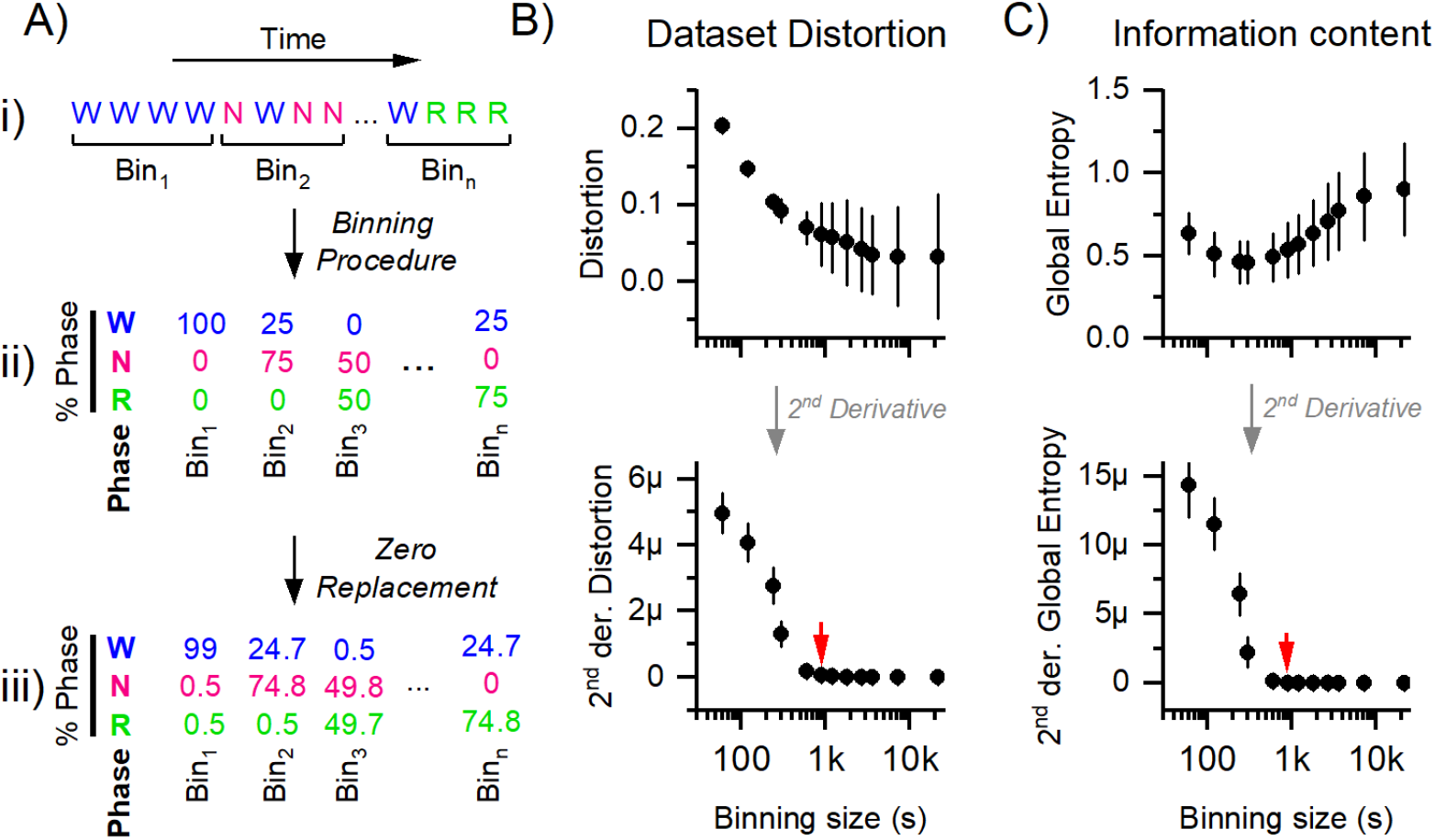
Optimization of binning window based on dataset distortion and information content. (A) Overview of the binning and zero-replacement procedures. (i) Example of sleep stage data across time, grouped into time bins (Bin_1_, Bin_2_, …, Bin_n_). (ii) For each bin, the percentage of time spent in Wake (W), NREM (N), and REM (R) is calculated. (iii) Zero-replacement is applied to address bins where one or more phases are undetected, substituting zero values (e.g., 0%) with small values (two-thirds of the detection limit) to enable compositional data transformation. (B) Quantification of distortion introduced by the zero-replacement procedure across varying bin sizes. Top: distortion between original and zero-replaced datasets decreases with larger bin sizes. Bottom: Second derivative of the distortion curve, with the inflection point (red arrow) indicating the optimal tradeoff between minimizing distortion and retaining resolution (∼900 seconds). (C) Evaluation of information content as measured by global entropy across binning windows. Top: Global entropy decreases with smaller bin sizes, indicating higher resolution but more noise; entropy increases at larger bin sizes due to oversmoothing. Bottom: Second derivative of the entropy curve, with the elbow point (red arrow) indicating the bin size where further aggregation yields diminishing gains in entropy (∼900 seconds).

As shown in Suppl. Fig. 1A, smaller binning windows resulted in a higher proportion of zero elements, as expected. Among the available zero-replacement methods^17^, we chose the computationally efficient approach of replacing zero elements with two-thirds of the detection limit (see Materials and Methods). To quantify the distortion introduced by this replacement, we computed the Hellinger distance between binned datasets with and without zero elements (Suppl. Fig. 1B). The Hellinger distance ranges from 0 (identical distributions) to 1 (maximally dissimilar) and served as a sensitivity metric for assessing the distortion caused by zero-replacement across bin sizes. As zero-replacement only affects zero elements, distortion was inversely proportional to the binning window size (Fig. 1B). We then identified the optimal binning window using the inflection point (or “elbow”) of the second derivative (Fig. 1B, red arrow, 900s), where additional binning minimally reduced the distortion at the cost of losing resolution.

We also examined entropy patterns across bin sizes. Larger bin sizes reduced noise at the cost of information, while smaller bin sizes retained information but increased noise. Entropy patterns across binning windows showed that smaller bins lowered entropy by capturing finer details, while larger bins raised entropy due to reduced precision (Fig. 1C). Again, we applied the second derivative method to identify the inflection point (Fig. 1C, red arrow), which also occurred at 900 seconds, indicating a tradeoff between information retention and noise reduction.

In summary, the choice of binning window can be informed by biological relevance, data-driven optimization, or a combination of both. Here, we introduced a principled approach based on entropy optimization and minimization of distortion from zero-replacement to guide binning in sleep architecture datasets.

### Compositional Data, Visualization and data analysis

Regardless of the chosen binning window, sleep architecture is typically represented and analyzed as the fraction or percentage of time spent in each sleep phase (Fig. 2A; W = Wake, N = NREM, R = REM). However, the compositional nature of this data, where all components are parts of a whole constrained to sum to a constant, has been largely overlooked. This oversight is critical, as traditional statistical methods are not suitable for compositional datasets and may yield biased or misleading interpretations^9^ (Suppl. Fig. 2).

**Figure 2:**
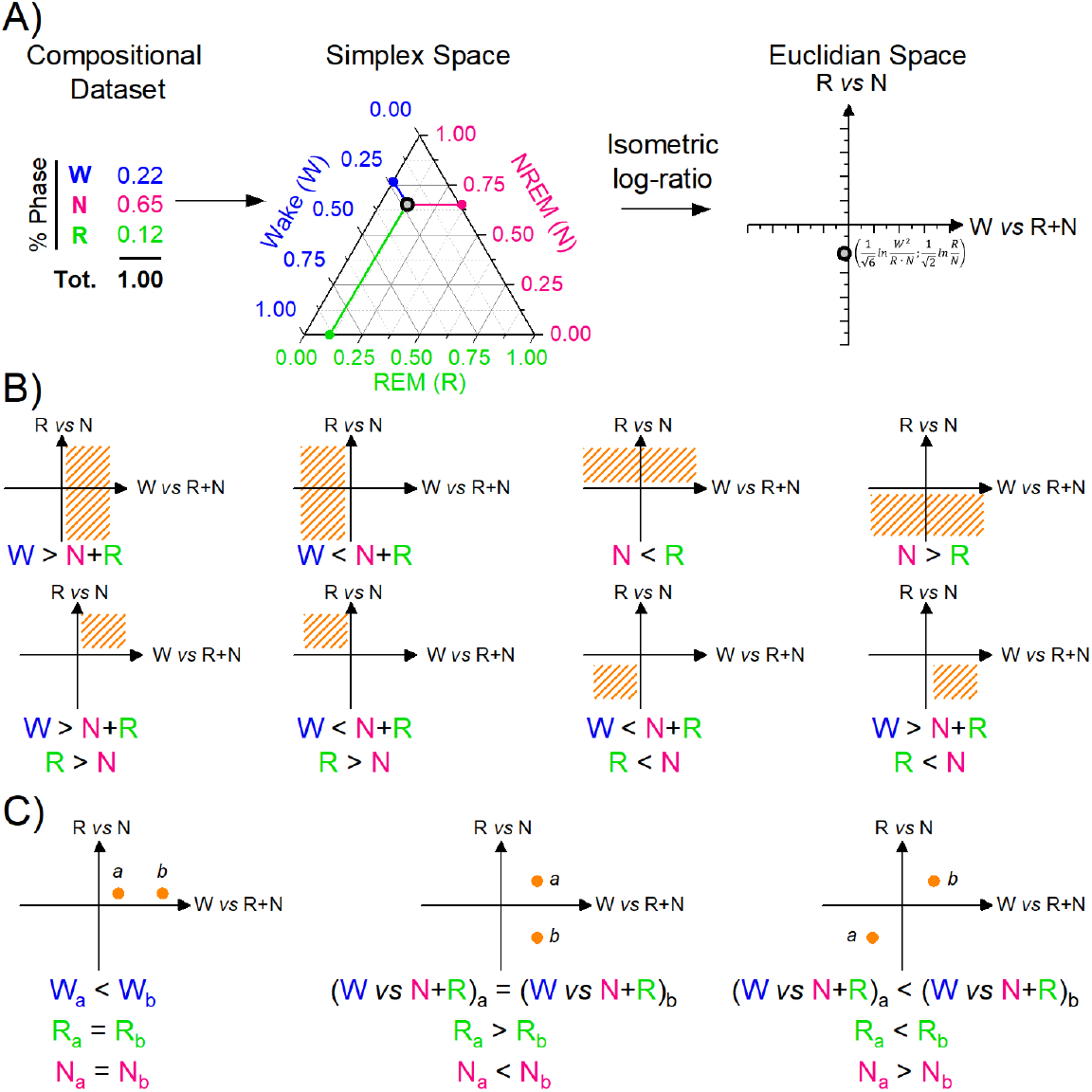
Transformation and interpretation of sleep architecture data using compositional and ILR frameworks. (A) An example of a compositional dataset representing the proportion of Wake (W), NREM (N), and REM (R) sleep phases. The data point is first represented in simplex space, constrained by the constant-sum property (W + N + R = 1), and then transformed into Euclidean space using the isometric log-ratio (ILR) transformation. The x-axis represents Wake vs. (NREM + REM), and the y-axis represents REM vs. NREM. Wake=W (blue); REM=R (green); NREM=N (magenta). (B) Schematic representation of possible positional outcomes in the ILR-transformed Euclidean space. (C) Interpretation of relative positions between two ILR-transformed data points (a and b).

To address this issue, we implemented an analytical strategy that respects the mathematical constraints of compositional data, while enabling accurate interpretation, appropriate statistical testing, and informative visualization. Our approach preserves the inherent interdependence among sleep phases, enhancing both the robustness and interpretability of the analyses.

Compositional datasets are characterized by non-negative components that collectively sum to a constant (typically 1 or 100%) and convey relative rather than absolute information. These datasets reside in a simplex space (Fig. 2A, Simplex space), which is a bounded, non-Euclidean geometry where standard operations (e.g., addition, subtraction) and assumptions (e.g., independence of components) do not hold ^9^. To perform valid statistical analysis, data must first be transformed from this simplex space into an unconstrained Euclidean space (Fig. 2A, Euclidean space), where conventional methods can be applied.

Several transformation techniques are available for this purpose, most of which rely on log-ratio transformations^18^. Among these, the isometric log-ratio (ILR) transformation offers two major advantages ^9^: it produces orthogonal (statistically uncorrelated) coordinates, and its components can be biologically interpreted. Applied to our three-part composition (Wake, NREM, REM), the ILR transformation yields two interpretable axes:

X-axis (Wake vs. Sleep): Represents the balance between wake and total sleep (i.e., Wake compared to the amalgamated NREM + REM phases).

Y-axis (REM vs. NREM): Represents the balance between REM and NREM sleep phases.

In this transformed Euclidean space, each quadrant can be interpreted based on the relative dominance of the sleep phases (Fig. 2B).

- Positive x-axis values indicate a dominance of Wake over Sleep;
- Negative x-axis values indicate Sleep dominance over Wake;
- Values near zero (x-axis) indicate a balance between Wake and Sleep;
- Positive y-axis values reflect a dominance of REM over NREM,
- Negative y-axis values reflect NREM dominance over REM, and
- Values near zero (y-axis) suggest equilibrium between REM and NREM.

This transformation allows for statistically valid and biologically meaningful comparisons, ensuring that the compositional structure of the data is respected throughout the analysis.

The relative position of two points in the ILR-transformed Euclidean space can be interpreted as shown in Fig. 2C. When two points (e.g., point *a* and point *b*) share the same y-axis value, they exhibit identical REM-to-NREM ratios. In this scenario, a point with a higher x-axis value (e.g., point *b*) indicates a greater dominance of wakefulness relative to total sleep (NREM + REM) compared to the other point. Conversely, if two points share the same x-axis value, their overall wake-to-sleep ratio is equal. However, differences in their y-axis values reveal shifts in the composition of sleep: a lower y-axis value (e.g., point *b*) indicates a greater dominance of NREM over REM. In this case, although wake is proportionally equivalent in both points, its interpretation is conditional on the internal balance between NREM and REM and cannot be interpreted in absolute terms between data points (i.e., we cannot comment about the absolute values of wakefulness between the two points, only relatively to total sleep). When two points differ on both axes, interpretation must account for both dimensions. A point with a higher x-axis value reflects increased dominance of wake relative to sleep, while a higher y-axis value indicates greater dominance of REM over NREM. Importantly, in all but the first scenario (equal y-axis), the absolute dominance of wake cannot be inferred in isolation; instead, wake must be interpreted relative to the combined sleep components, due to the compositional constraints.

After applying the zero-replacement procedure and performing the ILR transformation, the resulting dataset was both biologically interpretable and compatible with conventional statistical tests. Figure 3 illustrates this process using two representative sleep trajectories. Initially, sleep composition is displayed in the simplex space (Fig. 3A), followed by its transformation into Euclidean space via ILR, shown for each binning window (Fig. 3B) and color-coded by average values at each zeitgeber time (ZT) (Fig. 3C).

**Figure 3:**
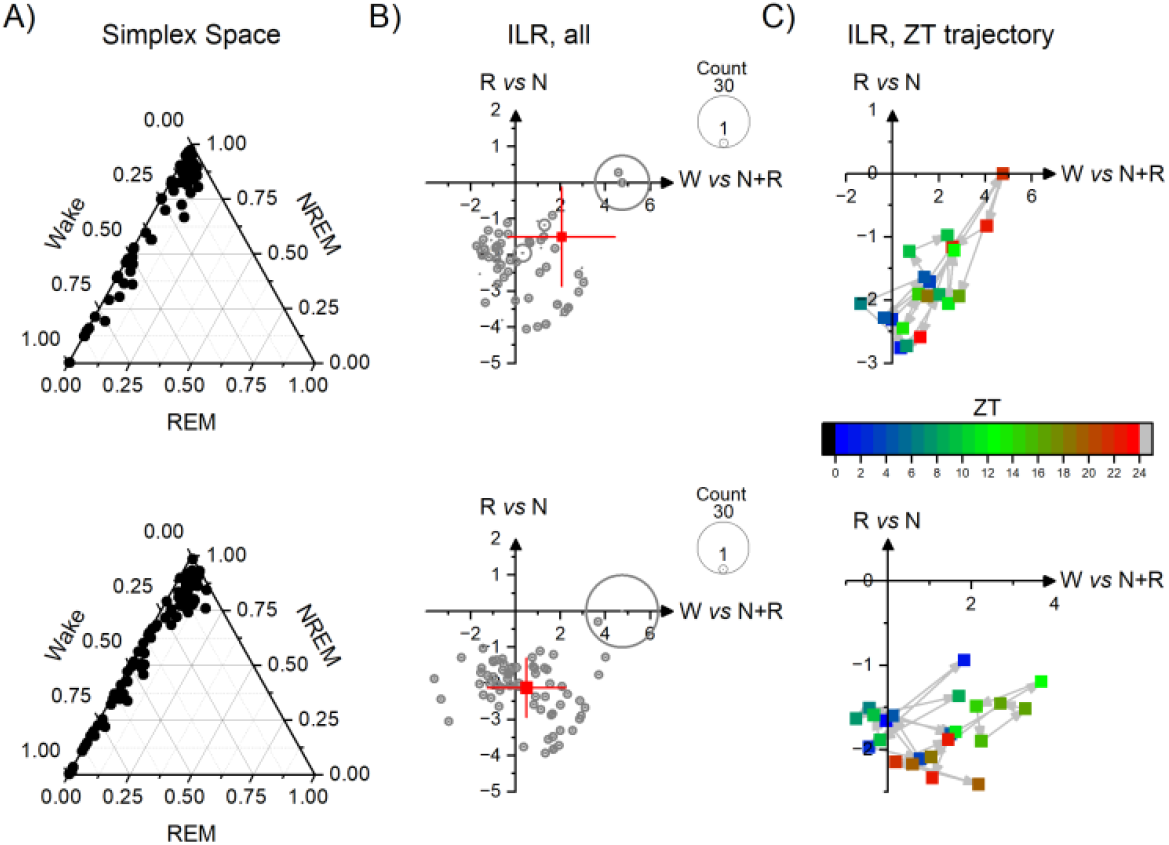
Visualization using compositional and ILR-transformed data. (A) raw data plotted in simplex space, illustrating how the sleep composition (Wake, NREM, REM) for each time point lies within the triangular boundaries defined by the constant-sum constraint. Both examples show the dominance of Wake, with most points clustered toward the Wake vertex. (B) corresponding data after ILR transformation, enabling projection into Euclidean space. The gray dots represent all time bins, with point size proportional to frequency. The red cross marks the center of mass (± SD) of the trajectory. In the top example, the cluster is shifted toward the positive x-axis, indicating a dominance of Wake over Sleep (Wake vs NREM+REM), and also slightly negative on the y-axis, reflecting a greater proportion of NREM over REM. The bottom example shows a similar Wake-dominant pattern but with a slightly broader distribution and a more central y-axis position, suggesting a more balanced NREM–REM ratio across time. (C) displays the ZT trajectory of ILR-transformed values, color-coded by zeitgeber time (ZT 0–24). In both examples, the ILR coordinates show temporal progression, the top trajectory shows a clearer shift between sleep and wake phases, while the bottom trajectory remains more variable and dispersed, suggesting a more fragmented or heterogeneous sleep pattern.

In Fig. 3B, the size of each point reflects the frequency of that specific stage configuration within the dataset, while the red marker denotes the center of mass (± standard deviation) of the full 24-hour sleep trajectory (ZT0 to ZT23), calculated by averaging the ILR coordinates across time. This transformation facilitates visual interpretation of the day-long sleep composition while sacrificing temporal resolution.

Further, the ILR-transformed data can be visualized in three dimensions, where the z-axis represents time (Suppl. Fig. 3). Figure 3C shows the 2D projection of this 3D trajectory onto the x-y plane, with the grey dot in Suppl. Fig. 2 corresponding to the points in the Euclidean Space.

In conclusion, the ILR transformation provides a powerful approach for converting compositional sleep data into biologically meaningful and statistically tractable variables, enabling robust visualization and analysis of sleep architecture over time.

### Transformed data, statistical approaches and considerations

Following ILR transformation, the dataset included two dependent variables for each subject, ilr_1_ (Wake vs. Sleep [NREM + REM]) and ilr_2_ (REM vs. NREM), measured across multiple time points (ZT), along with grouping information such as genotype (Fig. 4, top). Given the multivariate structure of the data (two dependent variables) and its repeated measures design (longitudinal data collected across time), we applied a one-way repeated measures MANOVA. This approach accounts for within-subject correlations over time while testing for group-level differences in global sleep architecture. If additional independent variables (e.g., sex, treatment) are of interest, the model can be extended to an N-way repeated measures MANOVA. In addition to the theoretical justification for applying a multivariate approach, and despite the expectation that the ilr transformation removes linear dependencies among coordinates, a repeated measures correlation (using *rmcorr* in R) revealed a significant residual correlation between the ilr-transformed coordinates (r=0.71, p<4·10^−77^, 95% confidence interval: [0.67, 0.76]). This further justifies the use of a multivariate modelling approach. Residual correlation can be explained by systematic, within-individual shifts in sleep architecture driven by factors such as circadian timing, sleep pressure, or environmental conditions, which consistently affect multiple sleep stages simultaneously.

**Figure 4:**
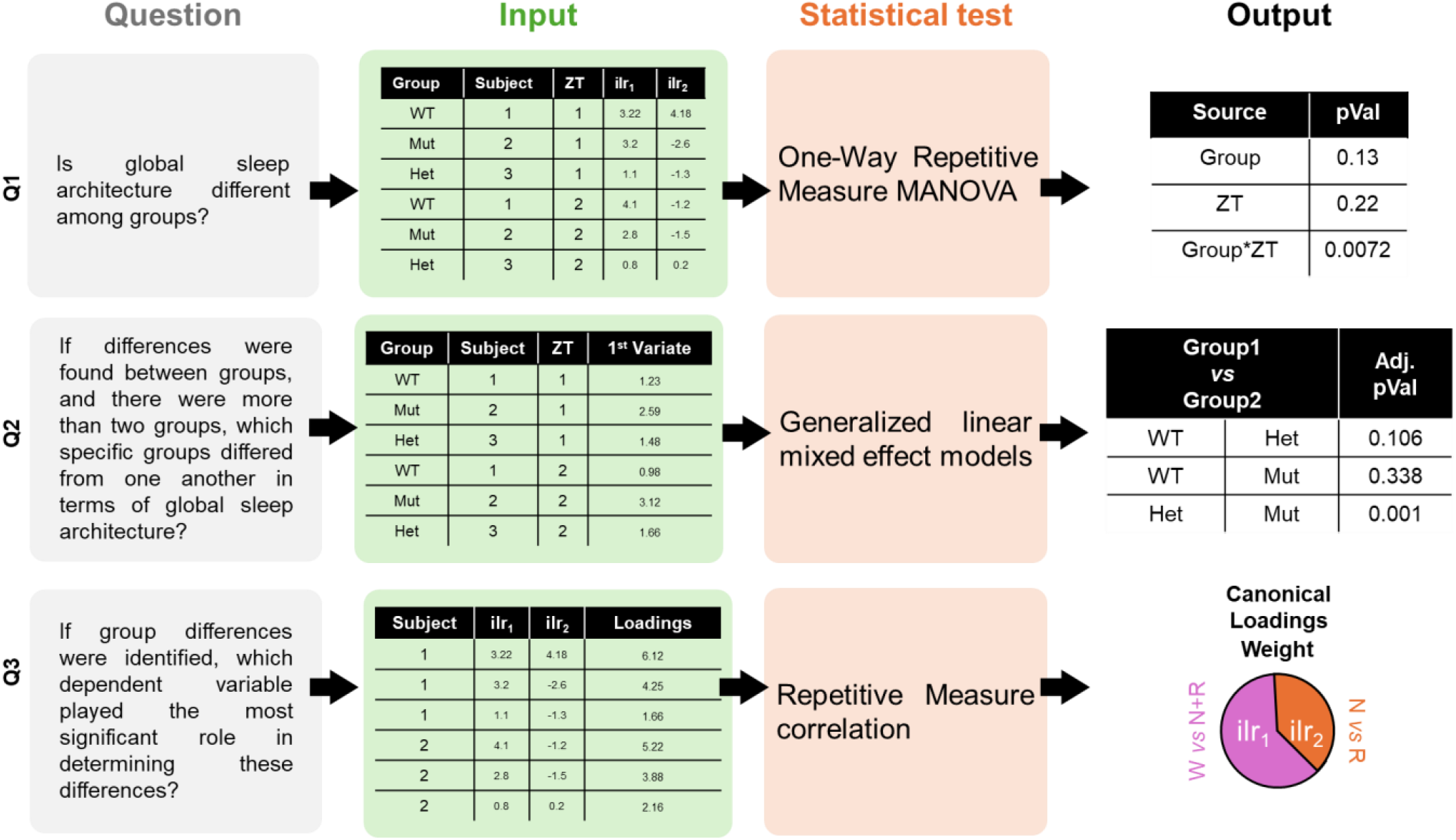
Statistical framework for analyzing ILR-transformed compositional sleep data. The diagram outlines a stepwise approach for evaluating group differences in global sleep architecture using ILR-transformed variables. Three key questions guide the analysis: (1) Are there group differences in sleep architecture? (2) Which groups differ? (3) Which ILR variable contributes most? A one-way repeated measures MANOVA tests global group differences. If significant, generalized linear mixed-effects models assess pairwise contrasts. Finally, repeated measures correlation and canonical loadings identify the most influential ILR variable (ilrq: Wake vs. Sleep, ilrq: REM vs. NREM).

This statistical framework enabled us to address several biologically meaningful questions: whether overall sleep architecture differed among groups (Q1); if more than two groups were included, which specific group pairs showed significant differences (Q2); and which variable, ilr_1_ or ilr_2_, contributed most strongly to group separation (Q3). The MANOVA tested for significant effects based on the combination of both dependent variables (referred to as variates) and also allowed for the evaluation of temporal dynamics by including ZT as a repeated measure.

In cases involving more than two groups, post-hoc testing is often necessary to identify specific group differences. However, most commonly used statistical software (e.g., SPSS, R) lack built-in support for multivariate post-hoc comparisons within a repeated measures framework. Although univariate post-hoc tests are frequently applied in practice, they are not statistically appropriate in a multivariate context^19^. To address this limitation, we propose the use of the variate that explains the largest proportion of variance (see Supplementary Material) as a new dependent variable in a univariate generalized linear model (Fig. 4, middle). This approach maintains the multivariate and repeated measures structure of the data while focusing on the principal source of group-level variation. However, it assumes that the first variate adequately summarizes the key features of the dataset; any meaningful variation explained by secondary variates is not captured and may affect interpretation.

To further understand the contribution of each transformed variable to the observed group differences, we examined the canonical loadings, the coefficients used to construct the variates (Fig. 4, bottom). These loadings provide a quantitative measure of how strongly each dependent variable (ilr_1_ or ilr_2_) influenced group separation. In simpler terms, they reveal whether differences between groups were primarily driven by Wake vs. Sleep balance or by the REM vs. NREM ratio. A detailed procedure for extracting canonical loadings from repeated measures MANOVA is provided in the Supplementary Material.

In conclusion, ILR-transformed compositional data allow for robust analysis of sleep architecture while properly accounting for its multivariate and longitudinal nature. Repeated measures MANOVA, combined with post-hoc modelling using the dominant variate and interpretation of canonical loadings, enables biologically meaningful insights into how different factors influence global sleep architecture across groups.

### Extending the pipeline to human data

In the present manuscript, we focused on the analysis of sleep architecture in mice, traditionally staged into three phases (Wake, NREM, and REM). The same concepts and analytical approaches discussed can also be extended to human data, where sleep staging is divided into Wake, REM, NREM-N1 (light), NREM-N2 (stable), and NREM-N3 (deep). While the graphical presentation may be more challenging to interpret due to the increased number of stages, we propose the following ILR transformations (respecting a proper sequential binary partition; Table 1 summarize the partition order), which retain biological interpretability:

**Table 1.** Sequential Binary Partition (SBP) of a human sleep staging. The SBP defines a hierarchical grouping of components used to construct balances for the isometric log-ratio (ilr) transformation. Each row represents a partition step that splits the composition into two groups: one group coded as +1 (included on the numerator side of the balance), another coded as −1 (included on the denominator side), and parts not involved in that particular split coded as 0. These groupings determine how each ilr coordinate is computed as a log-contrast of the corresponding parts.

| Partition Order | Wake | REM | NREM -N1 | NREM-N2 | NREM-N3 | r | s |
| --- | --- | --- | --- | --- | --- | --- | --- |
| 1 – ILR1 | +1 | -1 | -1 | -1 | -1 | 1 | 4 |
| 2 – ILR2 | 0 | +1 | -1 | -1 | -1 | 1 | 3 |
| 3 – ILR3 | 0 | 0 | +1 | -1 | -1 | 1 | 2 |
| 4 – ILR4 | 0 | 0 | 0 | +1 | -1 | 1 | 1 |

- ILR1: Wake vs. sleep (REM + N1 + N2 + N3), relative proportion of wakefulness compared to all sleep stages.
- ILR2: REM vs. NREM (N1 + N2 + N3), relative proportion of REM compared to all NREM stages.
- ILR3: N1 vs. N2 + N3, relative proportion of light NREM (N1) compared to deeper NREM stages (stable N2 and deep N3).
- ILR4: N2 vs. N3, relative proportion of stable NREM (N2) compared to deep NREM (N3).

It is important noticing that the ILR transformation for the scheme presented in Table1, each ilr coordinate is computed as follows:

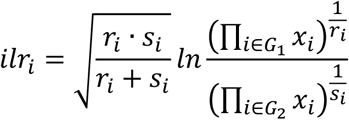

where *r_i_* is the count of +1 elements, *s_i_* is the count of -1 elements, *G*_1_ is the group of +1 elements and *G*_2_ is the group of -1 elements. In other words, this is equivalent to computing the log ratio of the geometric means of the two groups, scaled by a factor based on their sizes. After the ILR transformation, the same statistical approaches suggested above remain valid, with only an increased number of dependent variables.

## Discussion

This study presents a principled analytical framework for the evaluation of sleep architecture, based on the recognition of its compositional structure and the benefits of data-driven preprocessing. By addressing two often-overlooked methodological challenges, binning strategy and the statistical treatment of compositional data, we offer a reproducible and statistically coherent pipeline that enhances the interpretability and reliability of sleep analyses.

Traditionally, sleep data are binned into fixed intervals, commonly one-hour segments, based on convention or experimental convenience. However, these arbitrary divisions may obscure meaningful temporal patterns or introduce analytical bias. Our data-driven binning strategy identifies an empirically justified window that minimizes the impact of zero-replacement while retaining information-rich temporal variability. This approach provides an objective, scalable method that can be adapted across datasets, offering a more nuanced understanding of sleep dynamics and enhancing reproducibility in preclinical sleep research.

Sleep architecture data are inherently compositional^9,20^. Traditional statistical methods, which assume independence among variables and operate in Euclidean space, are incompatible with this structure and risk distorting findings. The ILR transformation resolves this conflict by mapping compositional data from the simplex space to an unconstrained Euclidean space, enabling the use of standard multivariate statistics while preserving the relationships between components^21^. Importantly, the ILR coordinates, Wake vs. Sleep and REM vs. NREM, offer intuitive, biologically meaningful contrasts that facilitate interpretation and visualization.

Log-ratio transformations require strictly positive input, necessitating the replacement of zero values^14^. We employed a computationally efficient method (substituting zeros with two-thirds of the detection limit) to enable transformation. While effective for datasets with <10% zero entries, this method introduces substantial distortion when zeros are more prevalent, as observed in our study ^9,14^. In such cases, more robust methods like multiplicative replacement (multRepl) may be used, although also these come with limitations. Notably, multRepl maintains the covariance structure among non-zero components but may distort the relationship between zero and non-zero elements^20^. Arguably, the preservation of the internal structure among non-zero elements is more conceptually important, particularly for high-dimensional biological data. To evaluate how different binning windows and replacement strategies affect compositional integrity, we used a data-driven approach to identify an optimal binning size. This involved minimizing the distortion introduced by zero replacement and maximizing information retention as measured by global entropy. However, we recognize that such data-driven binning can limit comparability across studies, especially when different bin sizes or zero-replacement strategies lead to different representations of the same underlying data.

In this manuscript, we prioritized conceptual clarity and accessibility, presenting a streamlined analytical pipeline that integrates compositional data transformation with standard statistical tools. We believe that widespread adoption of these principles will improve the robustness and reproducibility of sleep research. As the field becomes more familiar with compositional data analysis, we anticipate the development and application of more sophisticated methods tailored to the specific structure of sleep data. It is also important to highlight recent findings by Le and colleagues^3,4^, who approached sleep architecture analysis as both compositional and longitudinal in nature. While they employed a similar ilr transformation of the compositional data, they used the resulting coordinates as independent variables, in contrast to our approach, where they serve as dependent variables. Furthermore, their work introduced a novel Bayesian statistical framework^3^ designed to handle the multivariate and compositional structure of sleep stage data. Their contributions reflect a growing awareness of these methodological considerations and demonstrate how appropriately modeling sleep architecture can lead to new and meaningful research questions.

Finally, we emphasize that compositional analysis has implications for how research questions in sleep science are formulated. For example, investigating whether “wakefulness increases” under a certain condition is incomplete unless considered in relation to concurrent changes in REM and NREM sleep. Our approach not only introduces a statistically appropriate method for analyzing sleep architecture but also encourages researchers to rethink the framing of biological questions in a way that respects the compositional and multivariate nature of the underlying data.

## Authors Contribution

M.B. and A.B.: conceptualization, data analysis, writing and editing.

## Supplementary Figures

**Supplementary Figure 1:**
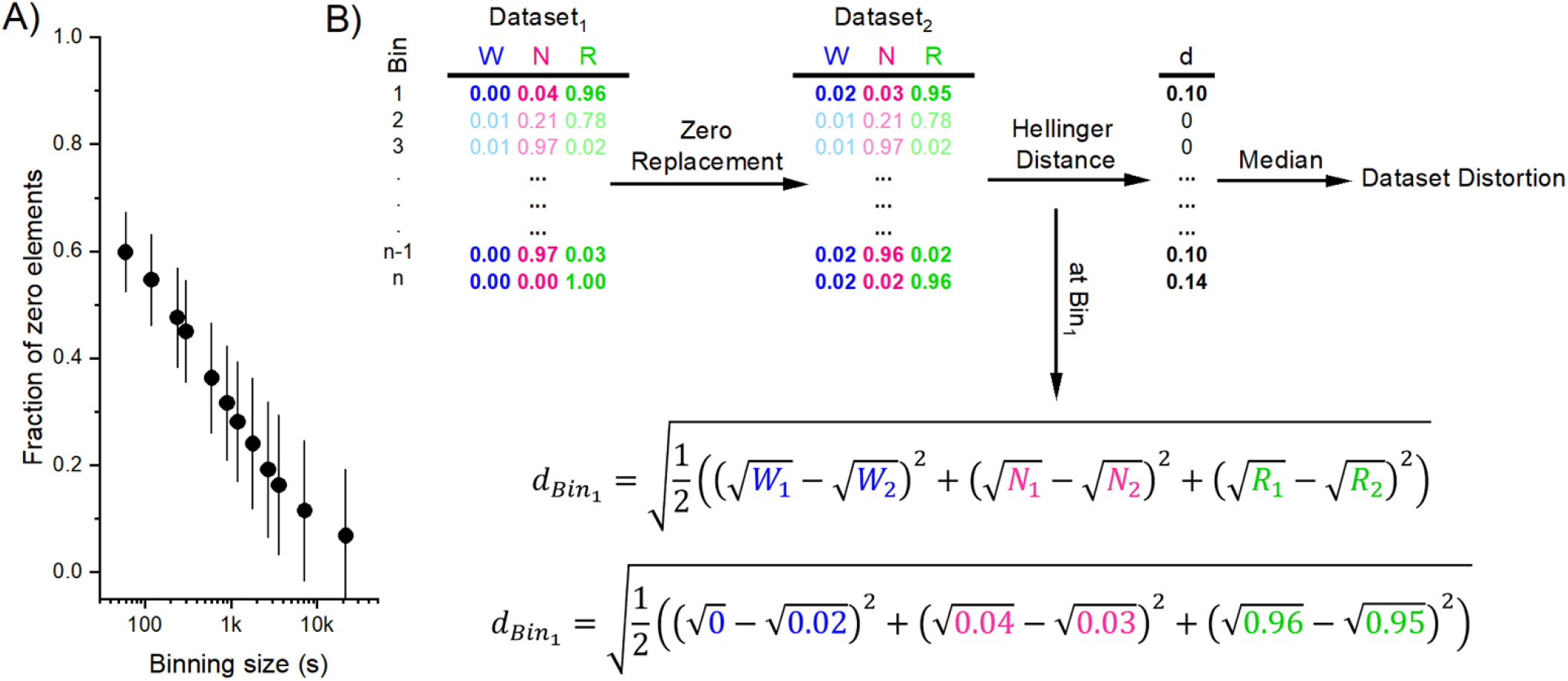
Impact of binning size on zero elements and dataset distortion in sleep architecture data. (A) Fraction of zero elements as a function of binning window size (in seconds). (B) Schematic representation of the distortion quantification process. A raw dataset containing zero elements (Dataset_1_) is subjected to zero-replacement, generating a new dataset (Dataset_2_) with small values substituted for zeros. The Hellinger distance is then calculated between the two datasets for each bin to quantify the distortion introduced by the zero-replacement procedure. The median of these distances across bins provides a global distortion metric. The lower portion of the panel shows the formula for Hellinger distance and its application using example values from Binq.

**Supplementary Figure 2:**
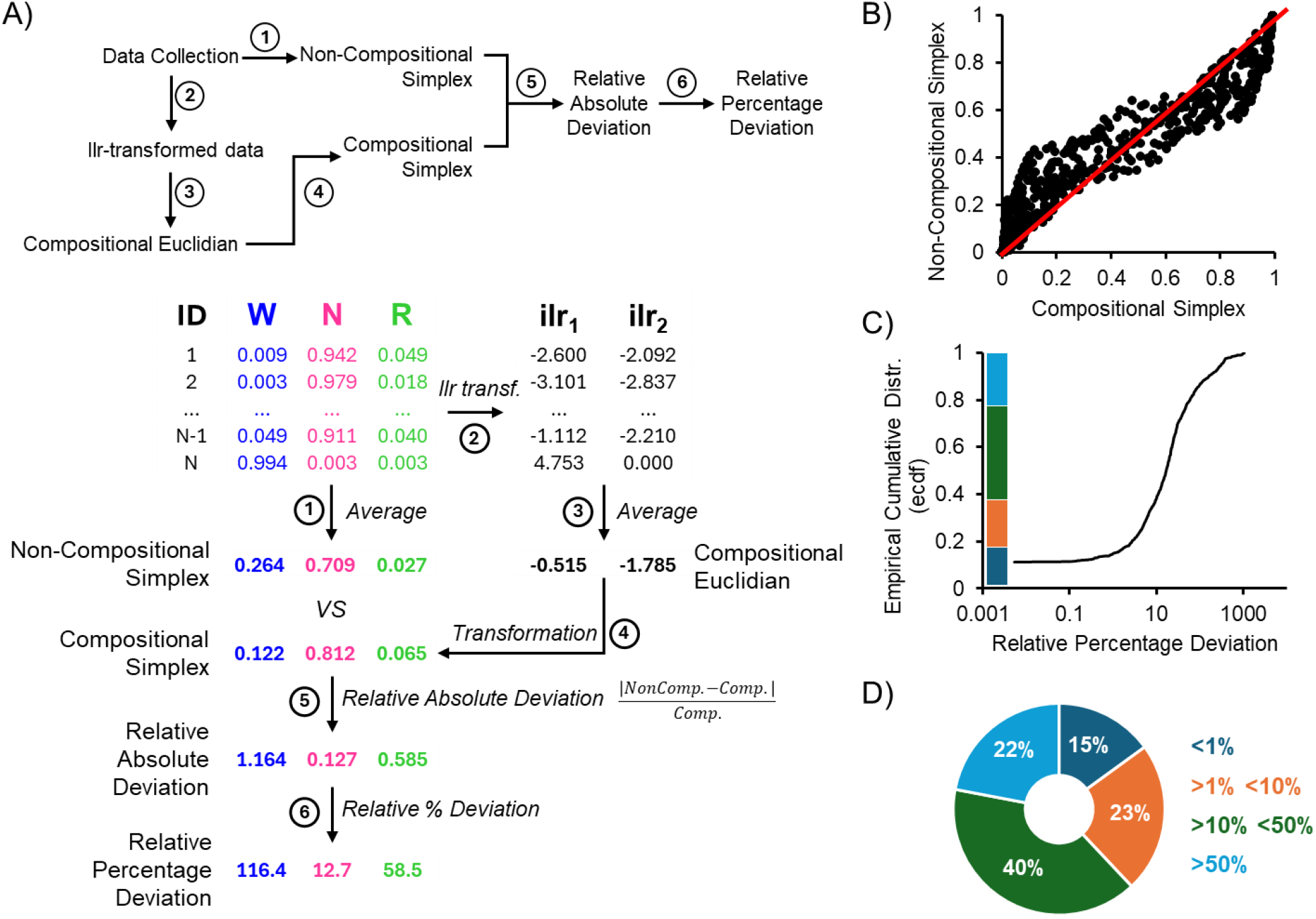
Impact of compositional versus non-compositional analysis on sleep architecture data. Transformation of sample data showing how compositional data (simplex-based) are preserved compared to distorted values when treated as non-compositional. (B) Scatter plot comparing values obtained through compositional (x-axis) and non-compositional (y-axis) methods. Deviations from the red line reflect systematic distortions. (C) Histogram showing the distribution of errors between compositional and non-compositional analyses, binned by error magnitude. (D) Pie chart summarizing the percentage of data points falling into each error range, indicating that 40% of points deviate by more than 10% when analyzed with non-compositional methods.

**Supplementary Figure 2:**
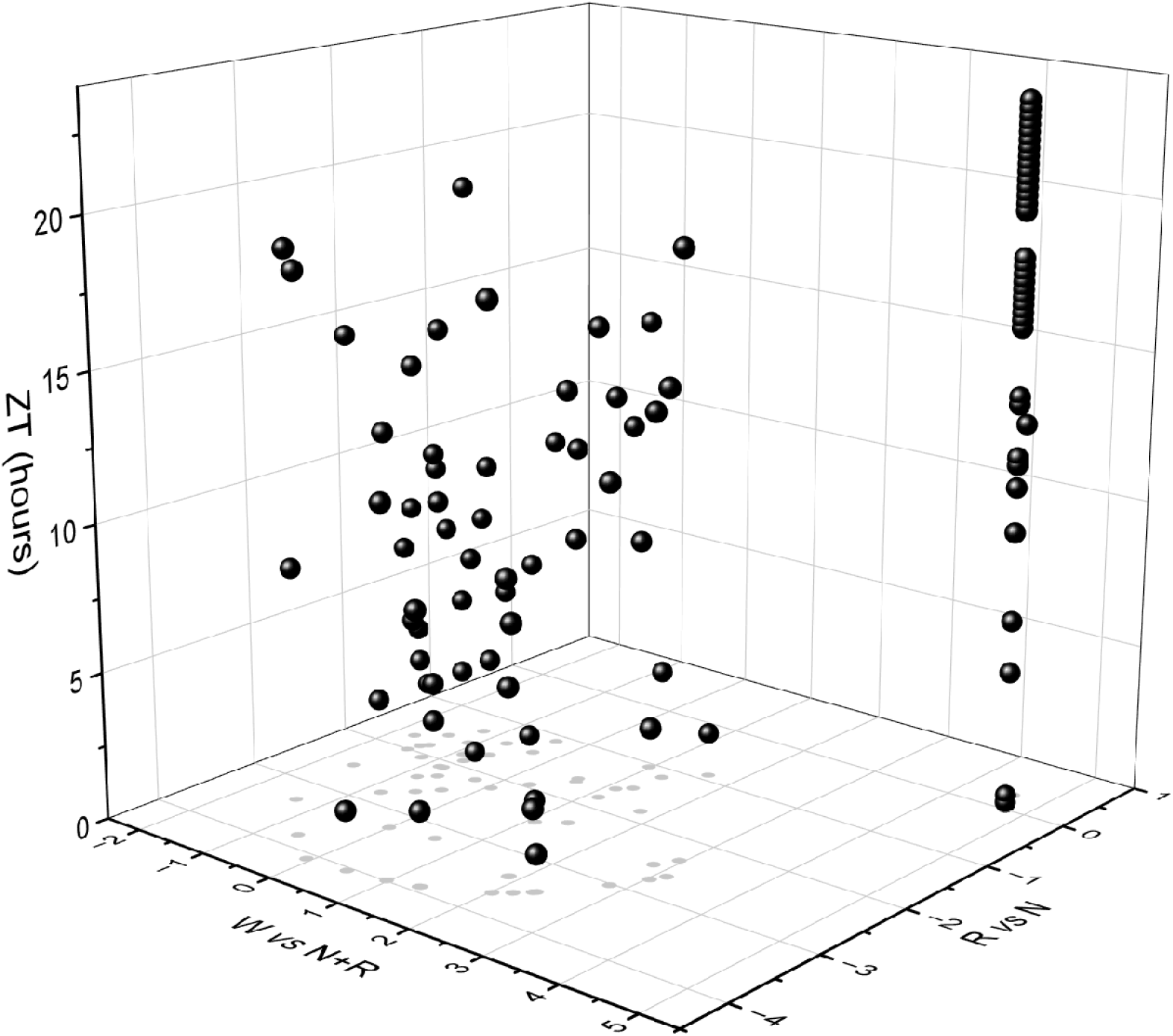
3D representation of sleep-wake data. ILR transformed data can be represented in space, adding changes across time on the Z plane.

## Supplementary Material

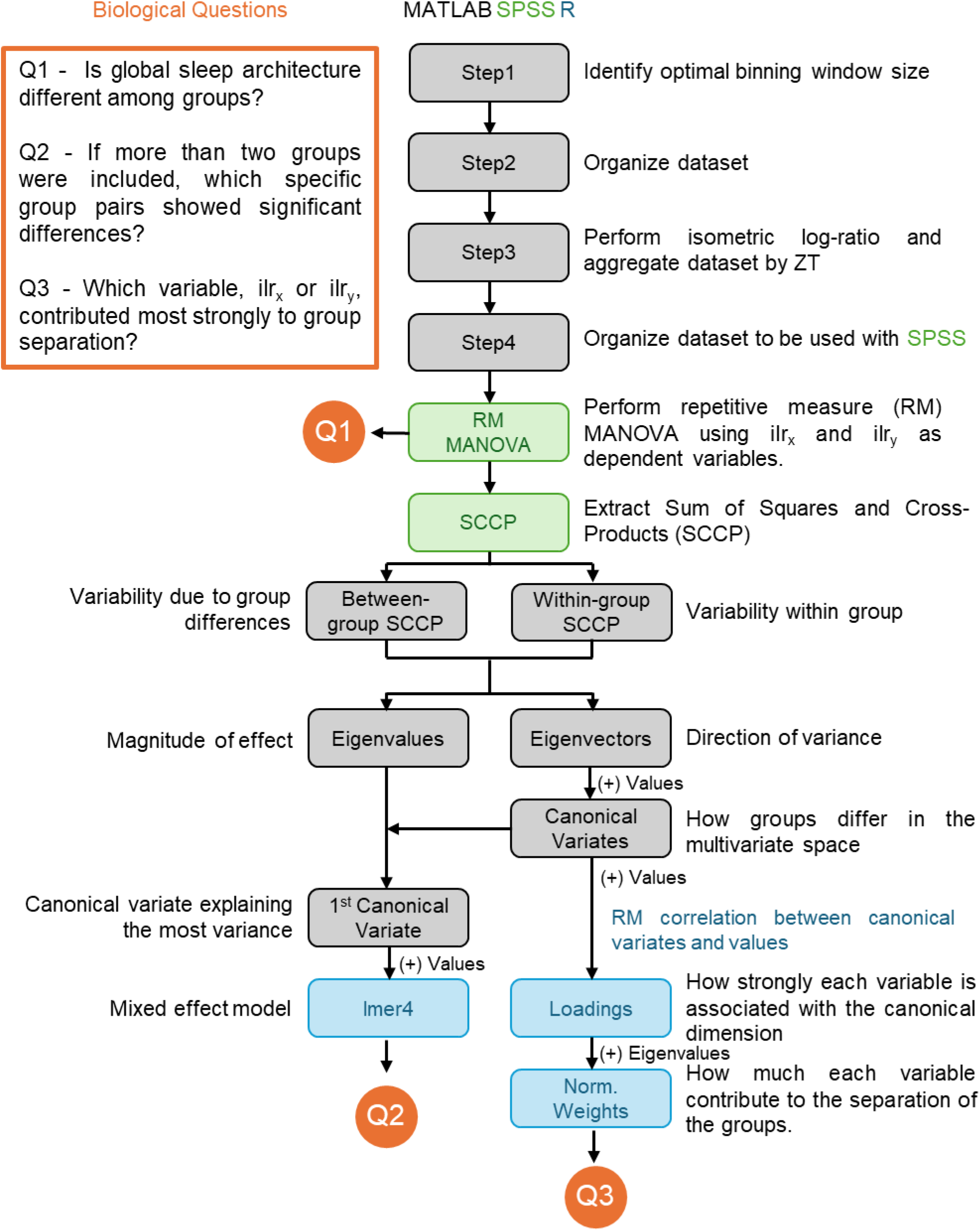

### Pipeline

Scripts and example files are found at https://doi.org/10.5281/zenodo.15707239.

### 1. Step1_FindBinSize.mat

<u>Aim</u> - to identify the optimal binning window size based on Hellinger distance and Shannon entropy.

<u>Short description</u> – the script is looping through all the text files present in the same folder and compute the Hellinger distance (Dataset Distortion) and Shannon entropy (Information Content) with varying binning window sizes. The aim is to identify (visually or by using the second derivative) the optimal binning window size based on the dataset.

<u>Requirements</u> – The script will run through the.txt files present in the same folder; accordingly, only the files to analyze in the.txt format should be present in the same folder with Step 1 script. A.txt file is used as input with at least the following columns: Time (mm/dd/yyyy HH:MM:SS format) and Episode (W, R, NR or N for wake, REM and NREM, respectively).

### 2. Step2_OrganizeDataset.mat

<u>Aim</u> - to create the binned dataset according to desired binning window size.

<u>Short description</u> – the script is looping through all the text files present in the same folder and compute the binned dataset to be used for further analysis.

<u>Requirements</u> – The script will run through the.txt files present in the same folder; accordingly, only the files to analyze in the.txt format should be present in the same folder with Step 1 script. A.txt file is used as input with at least the following columns: Time (mm/dd/yyyy HH:MM:SS format) and Episode (W, R, NR or N for wake, REM and NREM, respectively).

### 3. Step3_IsometricLogRatioOperations.mat

<u>Aim</u> – to perform the isometric log-ratio transformation on single binning windows and points aggregated at different ZT times.

<u>Short description</u> – For each binning window, the compositional dataset is transformed using the isometric log-ratio (ilr) method. After transformation, the x-axis represents the fraction of wakefulness compared to total sleep (REM + NREM), while the y-axis represents the fraction of REM compared to NREM sleep. The transformed data are then grouped by ZT, and finally, the sleep stage proportions are recomputed back into a compositional format.

<u>Requirements</u> – The Results from *Step2_OrganizeDataset.mat* needs to be computed first.

### 4. Step4_OrganizeDatasetForSPSS.mat

<u>Aim</u> – To organize the dataset to be used in Step5 and SPSS (Repetitive measure MANOVA)

<u>Short description</u> – The values are organized to be an adequate input for Repetitive measure MANOVA and Step5.

<u>Requirements</u> – The Results from *Step3_ IsometricLogRatioOperations.mat* needs to be computed first.

### 5. Repetitive measure MANOVA

To perform a repetitive measure MANOVA, we imported the Excel file generated from *Step4_OrganizeDatasetForSPSS.mat* into SPSS. To perform the analysis, select Analyze **→**General Linear Model **→**Repeated Measures, as shown below:

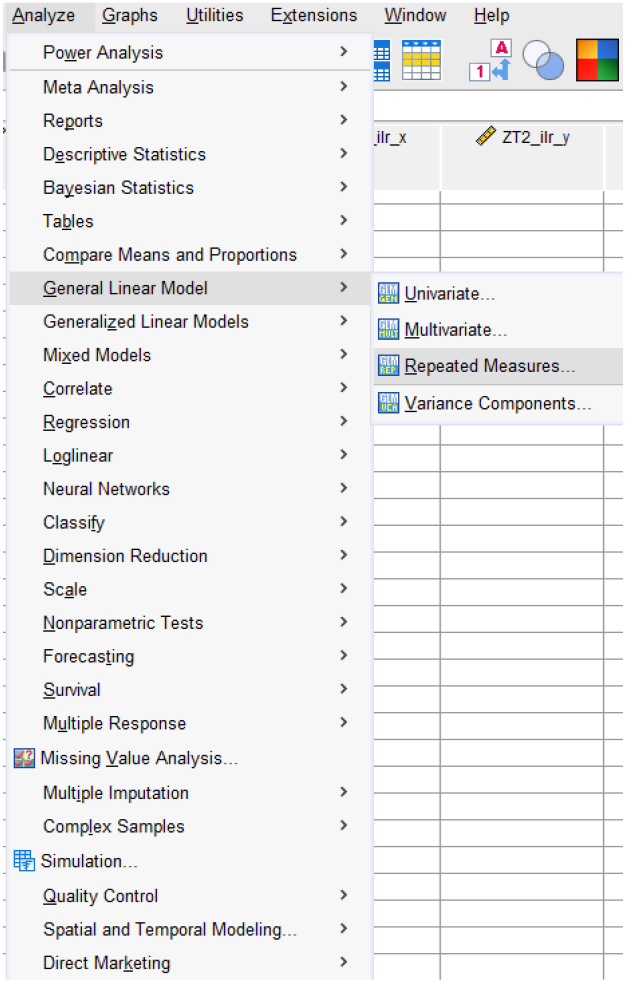

The following Menu window will appear:

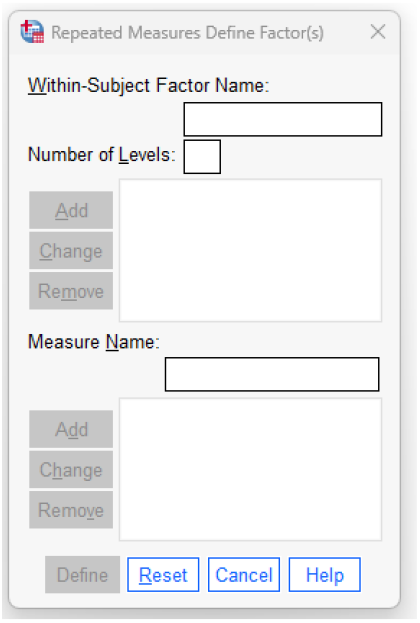

The within-subject factor in the dataset is ZT, and the two dependent variables are ilr_x (simplified in X) and ilr_y (simplified in Y). For the within-subject factor, the number of levels needs to be indicated, which are equivalent to the number of ZT values needed to be analyzed; for simplicity, a dataset with six ZT times was used. The resulting window will be:

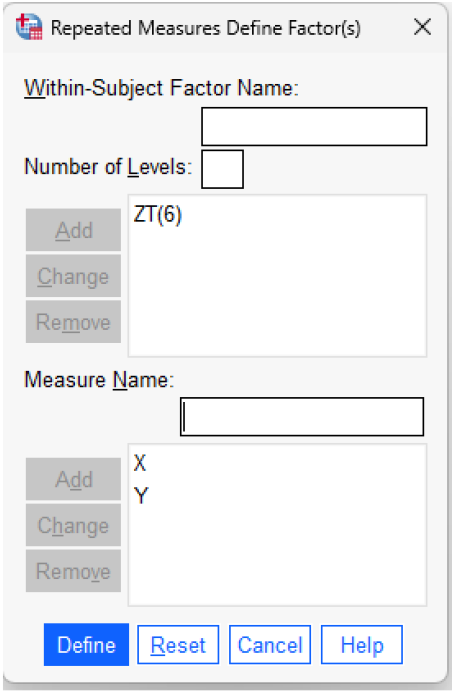

ZT(6) means that the software is expecting six levels of the within-subject factor, namely ZT. Given six levels of ZT for two dependent variables, the total number of columns of data are 12. In fact, each combination of within-subject levels and dependent variables need to be listed as separate columns (i.e., ZT1_ilr_x_, ZT1_ilr_y_, ZT2_ilr_x_, ZT2_ilr_y_, etc.). After clicking Define, the following window appears:

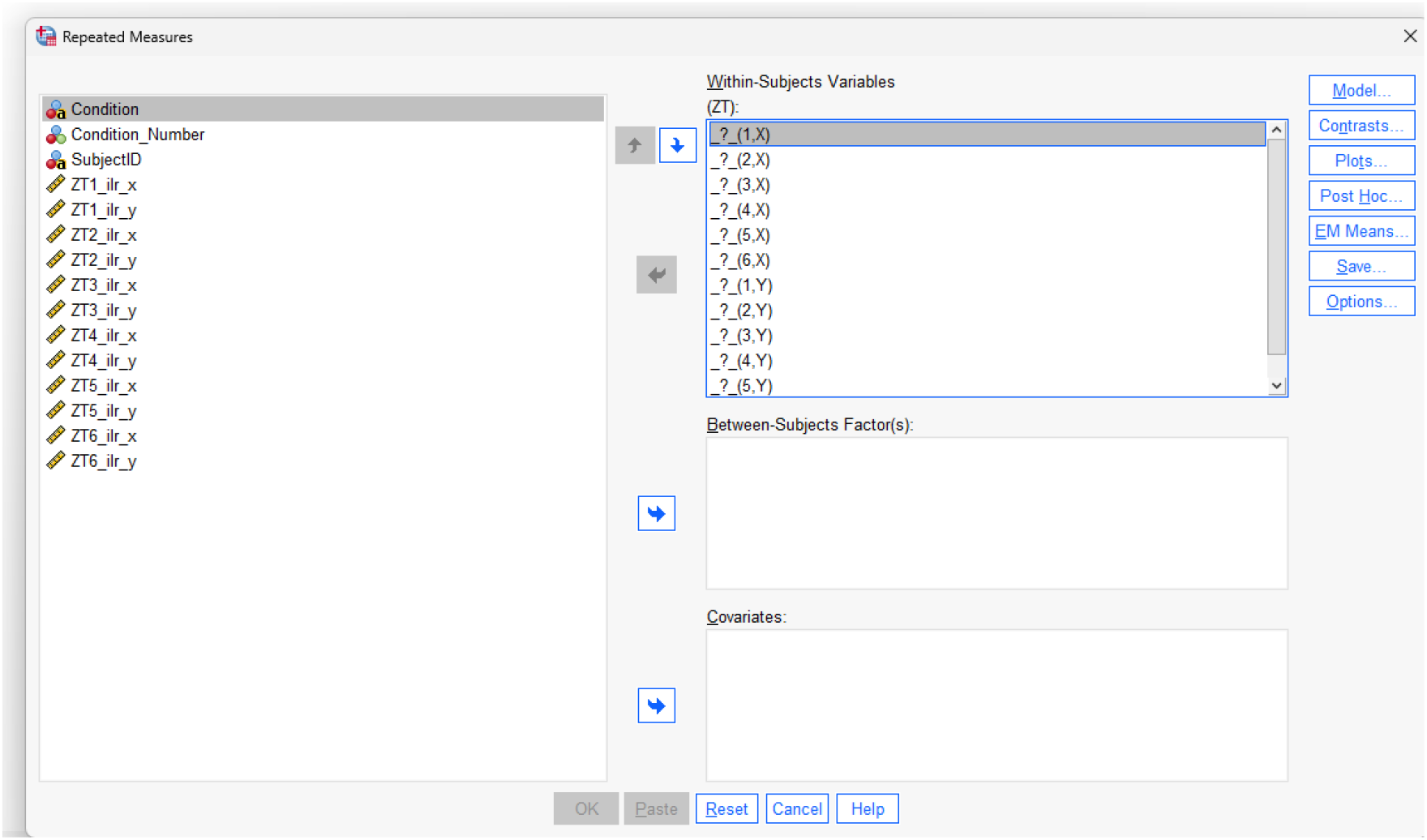

On the left side, there is a list of all the columns present in the dataset. Condition will be used as a between-subject factor. In our dataset, we did include a single between-factor, but more can be added (i.e., sex). In addition, while we did not include any covariate (i.e., age, weight of the mouse, etc.), they can be included in the Covariates section. On the Within-subjects variables window, we need to assign each column to the proper combination of within-subject levels (1 to 6 in the examples) and the dependent variables (X and Y in the example). Accordingly, _?_(1,X) is asking to assign the column with ZT1 and the X variable (ZT1_ilr_x). Following the same logic, we completed the selection as follows:

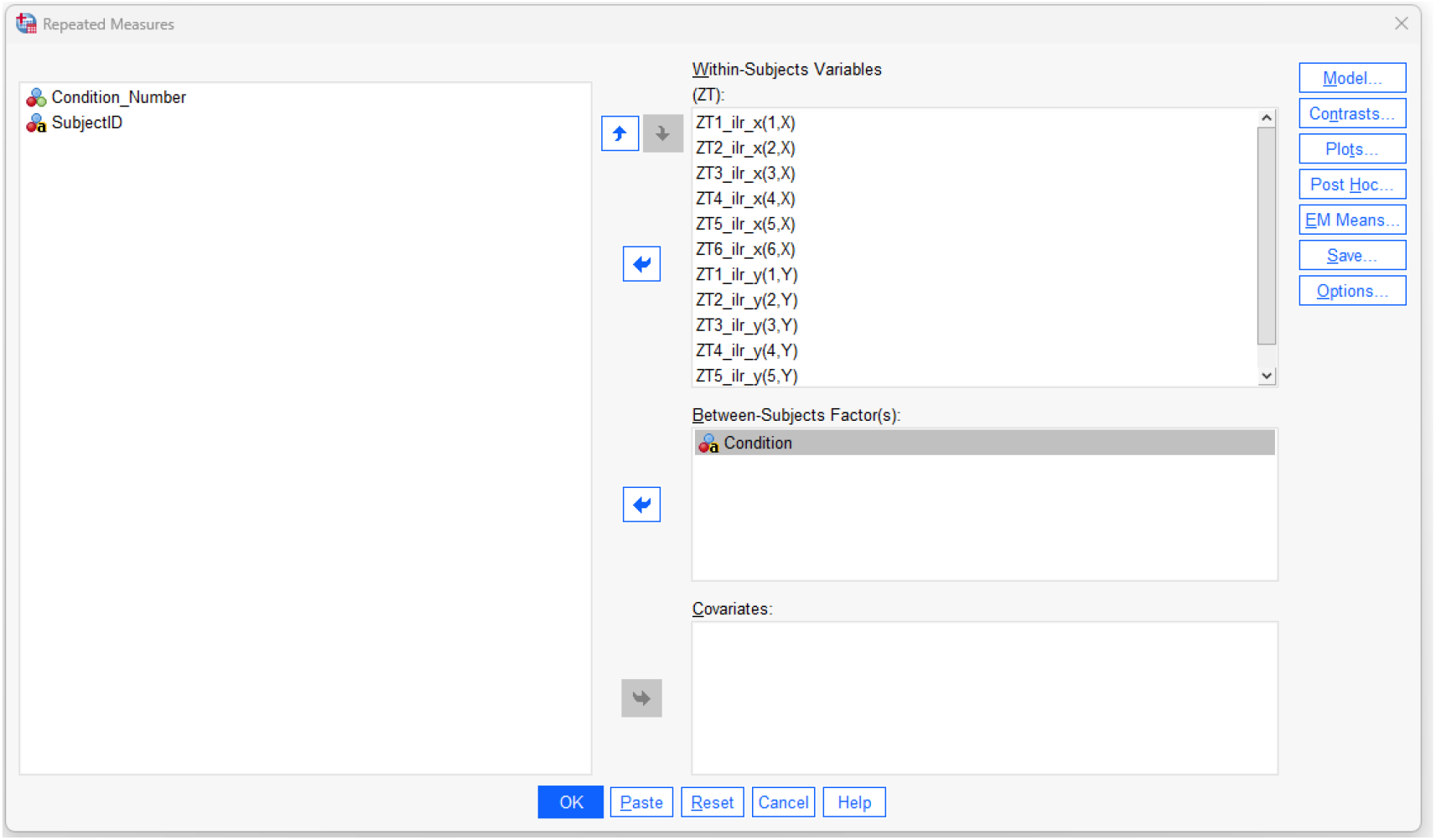

Finally, we need to be sure to select the SCCP matrix as an output; this can be done by selecting Options **→**SCCP matrices as shown in:

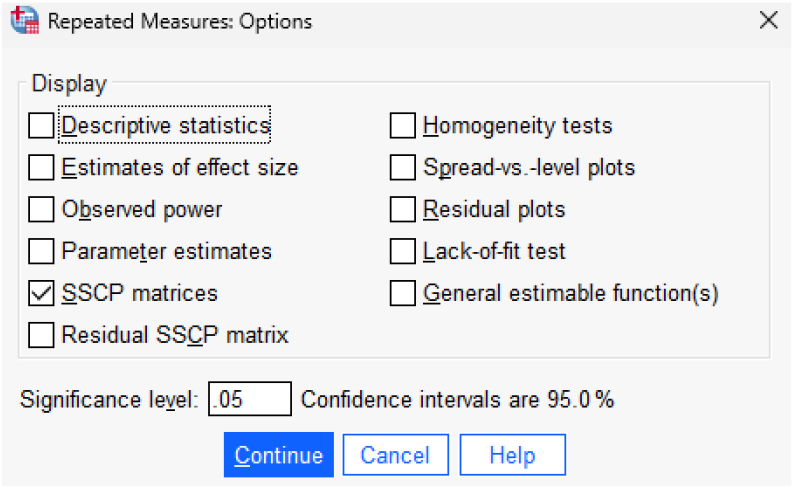

After this the model can be run and the output page appears.

The Multivariate Tests matrix can be used to answer Q1 (Is global sleep architecture different among groups?):

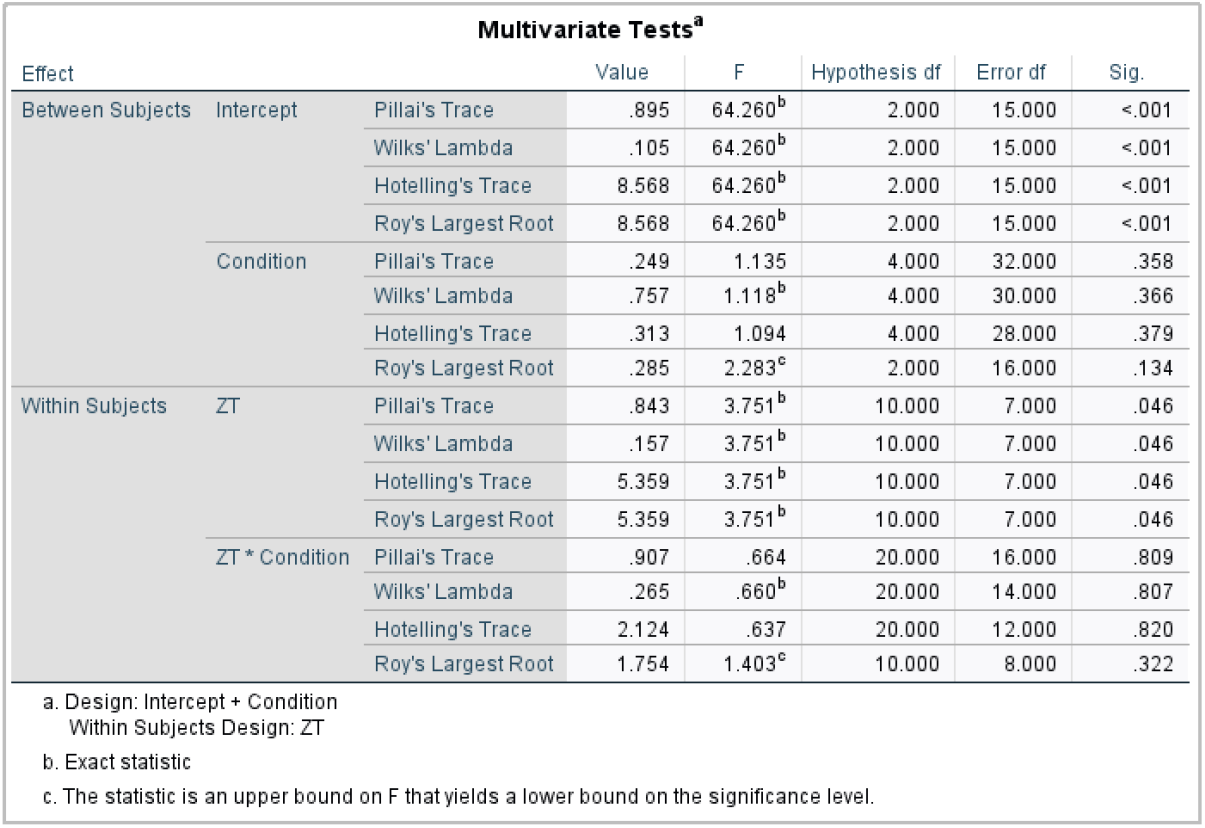

To interpret the results in this multivariate analysis table, it’s important to understand that the test evaluates the effects of both between-subjects (Condition in the example) and within-subjects factors (ZT in the example) on multiple dependent variables simultaneously (X and Y in the example). The four main multivariate test statistics—Pillai’s Trace, Wilks’ Lambda, Hotelling’s Trace, and Roy’s Largest Root —are used to assess whether these effects are statistically significant. These tests help avoid issues related to running multiple univariate tests while accounting for correlations among dependent variables. The choice of the test to apply is based on the study design, statistical power considerations, and assumptions such as sphericity or sample size; for instance, Pillai’s Trace is generally recommended when assumptions are violated, or sample sizes are small. The significance of the Intercept indicates that the overall mean of the dependent variables across all conditions and time points is significantly different from zero. The Condition effect (main effect; between-subjects factor) is not statistically significant, suggesting that there are no meaningful differences between the experimental groups and independently from ZT (within-subject factor) when considering all dependent variables together. While considering ZT (main effect; within-subject factor) independently from Condition (between-subject factor), the p-values for all tests are 0.046, indicating a significant effect of time independently on the Condition. Finally, when considering the changes of ZT based on Condition (interaction; ZT*Condition), no significance emerged, suggesting that the trajectories over time are similar in all Conditions.

From this analysis, we also extract the SCCP matrix (double click on it to copy the desired numbers):

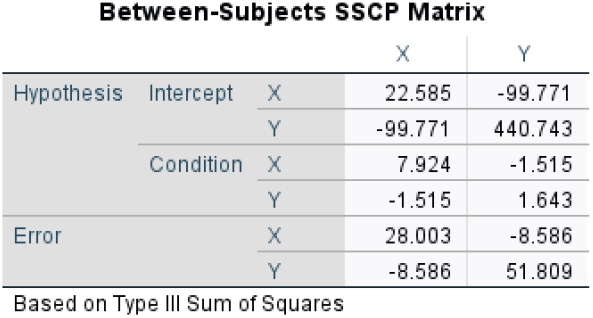

The SCCP (Sum of Squares and Cross-Products) is a symmetric matrix that summarizes the relationships between variables in a dataset. The matrix captures the variance of each variable along the diagonal and the covariance between pairs of variables in the off-diagonal elements (same numbers). The Intercept block captures the overall variability across all subjects. The Condition block captures the variability attributable to the between-subject factor (condition in this example). The Error block captures the variability not explained by the model. The SCCP matrix is typically decomposed into two components:

- Between-Subject Matrix (B) - captures variability due to group differences (Condition Block)
- Within-Subject Matrix (W) - captures variability within groups (Error Block)

To answer Q2 (if more than two groups were included, which specific group pairs showed significant differences?), and Q3 (which variable, ilr_x_ or ilr_y_, contributed most strongly to group separation?), we need to compute the canonical variates and canonical values, which can be computed by solving the inverse eigenvalue problem as follows:

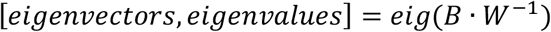

where eigenvectors represent the directions of maximum variance in the data and the eigenvalues measures the magnitude of the effect size for each eigenvector. *Step5_ComputeVariates.mat* is used to solve the inverse eigenvalue problem.

### 6. Step5_ComputeVariates.mat

<u>Aim</u> – To compute eigenvalues, eigenvectors and variates.

<u>Short description</u> – The between-subject and within-subject matrices computed from the SCCP matrix (repetitive measure MANOVA output from SPSS) are used to compute the eigenvalues, eigenvectors and canonical variates.

<u>Pseudocode</u>

#### Algorithm

Step5_ ComputeVariates

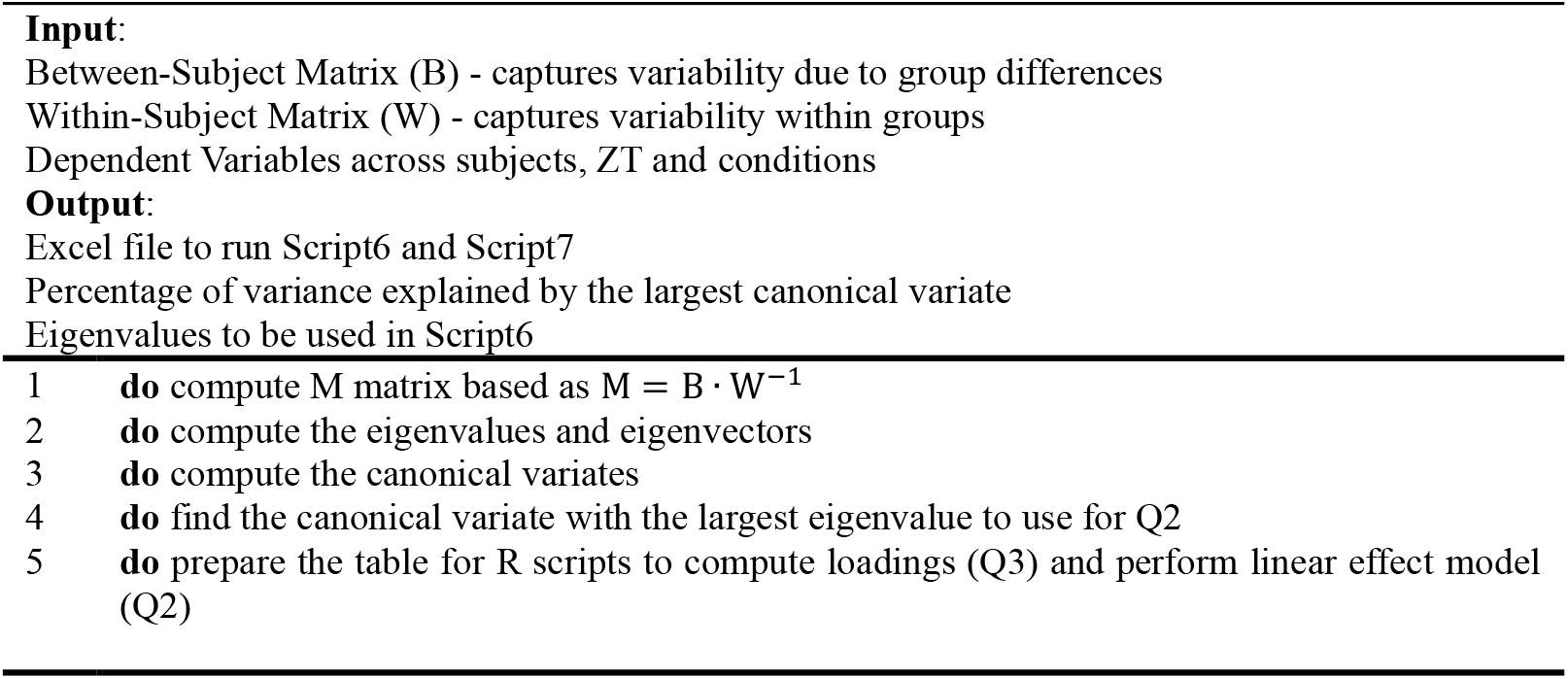

The canonical variates are a proxy of how the groups differ from each other in a multivariate space. To compute the canonical variates, we used the matrix with the ilr_x_ and ilr_y_ values (a *n* x *p* matrix, where *n* is the number of data points and *p* is the number of dependent variables) and the eigenvector matrix (a *p* x *k* matrix, where *p* is the number of dependent variables and *k* is the number of significant eigenvalues. First, we centered the data because the eigenvectors are derived from the covariance structure, which works under the assumption of mean-centered data. Then, we projected the centered data onto the eigenvectors.

The selection of the first covariate, to use as dependent variable in a mixed effect model, is done by selecting the variate with the largest eigenvalue. Since the robustness of the statistic is dependent on the explained variance by the first covariate, we printed the result in the command window of the script.

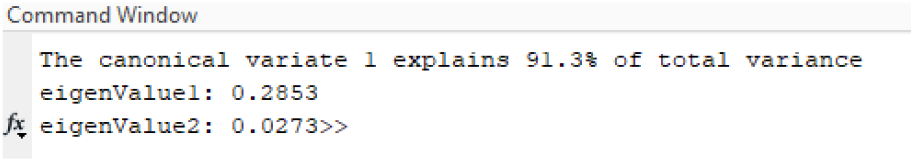

This percentage of variance explained by the largest covariate should be then properly reported in the Results section when answering Q3.

### 7. Step6_LoadingComputation.R

<u>Aim</u> – Compute the canonical loadings and determine which dependent variable weighted more in the group separation.

<u>Short description</u> – Using the original values (ilr_x_ and ilr_y_), canonical variates and eigenvalues, we can compute the percentage of importance of each dependent variable in separating the groups in the multivariate space.

<u>Pseudocode</u>

#### Algorithm

Step6_LoadingComputation

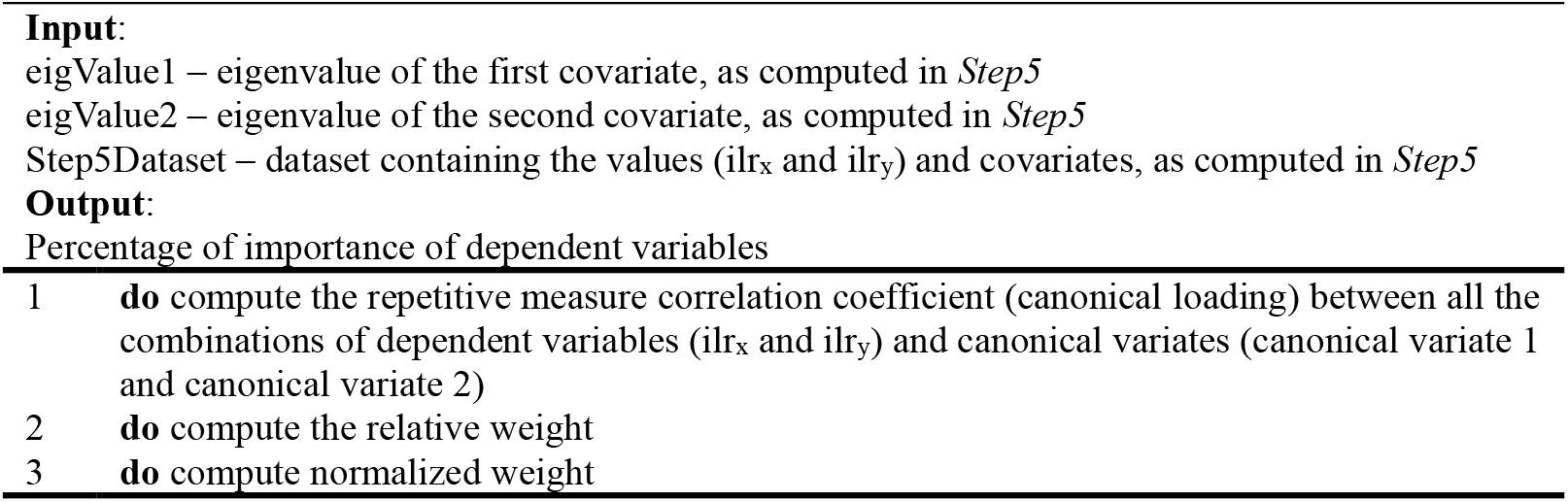

Canonical loadings are defined as the correlation between original variables (ilr_x_ and ilr_y_) and the canonical variates. They are a proxy of how strongly each variable is associated with the canonical dimension. Since our experimental design included also a repetitive measure component (ZT), we performed repetitive measure correlation in R. Higher is the canonical loading value, stronger that variable was associated with the canonical variate.

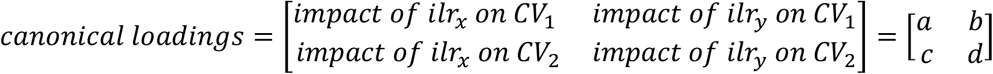

To compute how much each variable (ilr_x_ and ilr_y_) contributed to the separation of the group in the overall model, we computed the weight as follows:

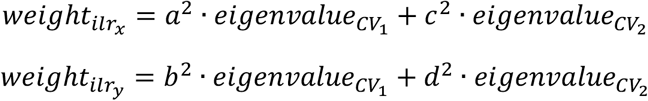

Finally, we normalize each value by the total weight carrying a final % value of importance:

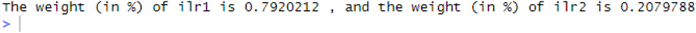

### 8. Step7_MixedEffectModel.R

<u>Aim</u> – To perform post-hoc analysis on groups

<u>Short description</u> – Using the canonical variate explaining the largest amount of variance (LargestVariate) as a proxy of the multivariate model, model the experimental design with a mixed-effect model.

Selecting the most appropriate mixed-effects model involves balancing theoretical justification, data structure, and model performance. While including more random effects improves realism, it can lead to convergence issues or singular fits—especially with limited data per subject. Thus, model selection should balance theory with feasibility, ensuring the random effects structure is supported by the data. In the example, the dataset contains one dependent variable (LargestVariate), a between-subject factor (Condition) and a within-subject factor (ZT). A good starting point is to model LargestVariate ∼ Condition*ZT+(1|SubjectID), which indicates that the intercept varies by subject. In other words, each subject is allowed to have its own baseline level of LargestVariate, accounting for individual differences while estimating the fixed effects of Condition and ZT. If convergence allows it, consider including random slopes, such as (ZT | Subject), which permit the effect of ZT to vary across subjects. In the script, it is also provided a post-hoc analysis (using false discovery rate to adjust for multiple comparisons) focused on comparing the conditions for a given ZT time.

